# Machine learning prediction of emesis and gastrointestinal state in ferrets

**DOI:** 10.1101/607242

**Authors:** Ameya C. Nanivadekar, Derek M. Miller, Stephanie Fulton, Liane Wong, John Ogren, Girish Chitnis, Bryan McLaughlin, Shuyan Zhai, Lee E. Fisher, Bill J. Yates, Charles C. Horn

## Abstract

Although electrogastrography (EGG) could be a critical tool in the diagnosis and treatment of patients with gastrointestinal (GI) disease, it remains under-utilized. The lack of spatial and temporal resolution using current EGG methods presents a significant roadblock to more widespread usage. Human and preclinical studies have shown that GI myoelectric electrodes can record signals containing significantly more information than can be derived abdominal surface electrodes. The current study sought to assess the efficacy of multi-electrode arrays, surgically implanted on the serosal surface of the GI tract, from gastric fundus to duodenum, in recording myoelectric signals. It also examines the potential for machine learning algorithms to predict functional states, such as retching and emesis, from GI signal features. Studies were performed using ferrets, a gold standard model for emesis testing. Our results include simultaneous recordings from up to six GI recording sites in both anesthetized and chronically implanted free-moving ferrets. Testing conditions to produce different gastric states included gastric distension, intragastric infusion of emetine (a prototypical emetic agent), and feeding. Despite the observed variability in GI signals, machine learning algorithms, including k nearest neighbors and support vector machines, were able to detect the state of the stomach with high overall accuracy (>80%). The present study is the first demonstration of machine learning algorithms to detect the physiological state of the stomach and onset of retching and could provide insight into methodologies to treat GI diseases and control symptoms such as nausea and vomiting.

## Introduction

Although electrogastrography (EGG) could be a critical tool in the diagnosis and treatment of patients with gastrointestinal (GI) disease, it remains under-utilized. The lack of spatial and temporal resolution using current methodologies to record GI myoelectric activity presents a significant roadblock to more widespread usage. The typical approach employs adhesive dermal electrodes -- similar to those used in electrocardiography -- placed on the abdomen, while the patient is instructed to remain still for baseline and postprandial testing [1]. These abdominal skin electrodes are not well-positioned to resolve functions of different GI compartments. Moreover, data collection is limited to brief artificial testing conditions, such as sitting in a chair to reduce motion artifacts. Implantable devices could provide greater resolution of GI signals as patients go about normal activities, including consuming meals, which are often associated with negative symptoms in patients with GI disease [2]. An implantable device could also serve as the input component for producing a closed-loop system to detect aberrant events, such as nausea in patients with gastroparesis, and to control therapeutic stimulation of the GI tract and its neural innervation [3].

Human and preclinical studies show that implanted GI myoelectric electrodes can record signals that contain significantly more information than could be derived from skin surface electrodes [4]. High-density electrode arrays placed on the human stomach during surgery provide insight into the directional propagation of signals in several compartments [5, 6]. Furthermore, animal studies are beginning to demonstrate the utility of implantable devices to record GI function, leading to proof-of-concept studies using miniaturized, wireless, and closed-loop device configurations [3]. Significant questions remain before these novel methods can be translated to the clinic, including (1) how many signals are needed to assess specific functions of the GI tract, (2) how to compensate for the intrinsic variability of GI anatomy and electrode placement between individuals, and (3) what features within the GI myoelectric signal are associated with functional changes in the GI system, such as nausea or digestion.

The current work focuses on addressing these issues by assessing the ability of implanted electrode arrays to record GI myoelectric signals from the serosal surface of the GI tract, and exploring the potential for achieving a personalized assessment of GI signals using machine learning algorithms to predict GI functional states, such as retching and emesis. Custom multi-contact conformal planar electrodes were placed on the serosal surface of the GI tract, including the gastric antrum, body, fundus, and the duodenum. Studies were performed using acute isoflurane-anesthetized as well as chronically implanted behaving ferrets. Testing conditions to produce different gastric states included gastric balloon distension, liquid diet consumption, intragastric infusion of emetine (a prototypical gastric emetic agent derived from syrup of ipecac [7]), and feeding. The ferret was used because it is the “gold-standard” model for emesis testing by industry; for example, in the development of 5-HT3 and NK1 receptor antagonists [8, 9]. Furthermore, an extensive database on vagus and gastric physiology is available for this species [e.g., 10, 11–16], and it is one of the few commonly-used animal models that possesses an emetic reflex, which is lacking in rodents and lagomorphs [17].

## Materials and Methods

### Animals

Experiments were performed on 10 adult purpose-bred influenza-free male ferrets (*Mustela putorius furo; Marshall BioResources, North Rose, NY, USA).* Animals were adapted to the animal facility for 31±17.5 (mean ± SD) days before surgery (see Table 1 for ages and body weights). GI myoelectric recordings were obtained from 7 animals prepared for acute experimentation. The other 3 ferrets were instrumented for chronic recordings. The University of Pittsburgh’s Institutional Animal Care and Use Committee approved all experimental procedures. These procedures conformed to the National Research Council *Guide for the Care and Use of Laboratory Animals* (National Academies Press, Washington, DC, 2011).

**Table 1:**
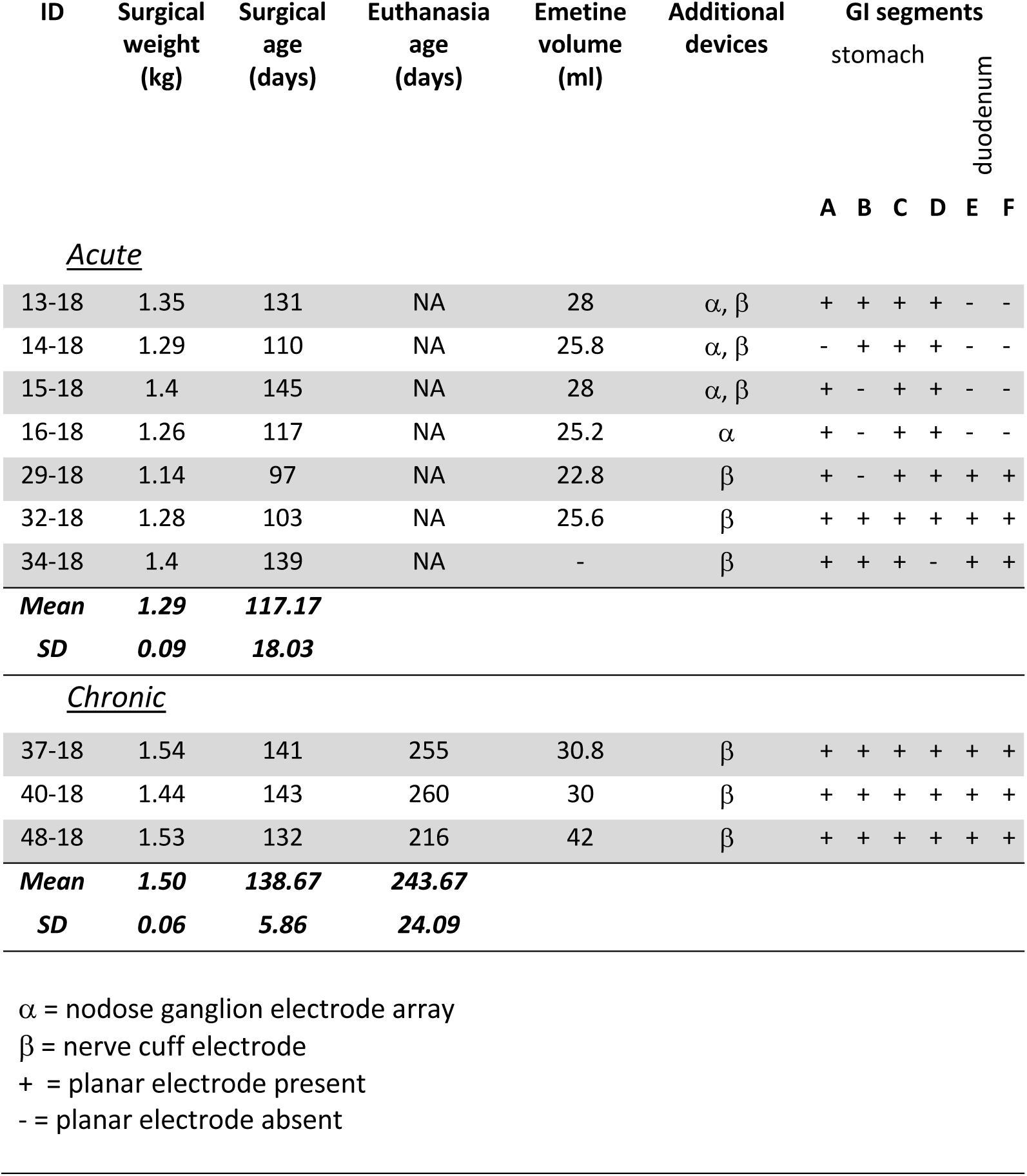
Subjects.

Animals were housed in wire cages (62 × 74 × 46 cm) under a 12 h standard light cycle (lights on at 0700 h), in a temperature (20-24°C) and humidity (30-70%) controlled environment. Food (ferret kibble; Mazuri Exotic Animal Nutrition, St. Louis, MI) and drinking water were freely available; however, food was removed 3 h before acute or chronic surgery and experimentation; all surgeries began at ∼0900 h. At the end of each study, ferrets were euthanized with an intracardiac injection of 1-5 ml euthanasia solution (390 mg/ml pentobarbital sodium; 5 mg/ml phenytoin sodium; SomnaSol EUTHANASIA-III Solution, Henry Schein Animal Health, Dublin, Ohio, USA) under isoflurane anesthesia (5%).

### Acute surgeries

Ferrets were anesthetized using isoflurane (5% induction, 1–3% maintenance) vaporized in O_2_. The level of anesthetic was adjusted to maintain areflexia (defined as no response to toe pinch), and stable heart and respiratory rates. Each animal was placed in the supine position, and the ventral surface was shaved and scrubbed with betadine. Rectal temperature was monitored and maintained between 36–40°C using either a warm water heating pad (Gaymar T/Pump) or an electric heating pad and an infrared lamp. EKG was monitored using alligator clips placed subcutaneously or clipped to the flanks, just below the axilla. A midline, vertical, 4-cm incision was made just above the trachea (below the level of the thyroid cartilage) and a tracheotomy was performed. After the tracheotomy, anesthesia was delivered through the intratracheal tube. Intratracheal airway pressure was monitored using an air pressure transducer (SAR-830/AP Small Animal Ventilator, CWE, Inc., Ardmore, Pennsylvania, USA), which was used to measure respiratory rate and the occurrence of emetic episodes [18]. Blood pressure was monitored and recorded using a fluid-filled catheter inserted through the left femoral artery (DBP1000 series Direct Blood Pressure System, Kent Scientific, Torrington, Connecticut, USA).

Animals underwent a laparotomy to expose the abdominal contents. For stomach distension, a customized pillow-type 30 ml barostat polyolefin balloon catheter (Ref # CT-BP-1017, Mui Scientific, Mississauga, Ontario, Canada) was advanced into the stomach through a ∼0.5 cm incision on the lateral edge of the gastric fundus. Additionally, an infusion catheter (silicone, ID .030” × OD .065” × Wall Thickness .0175”, Ref # 807000, A-M Systems, Carlsborg, Washington, USA) was advanced into the stomach through the same incision site; the catheter tip was advanced to rest in the gastric antrum. The deflated barostat balloon and the infusion catheter were secured in place by tying a purse-string suture at the incision site and applying medical-grade tissue adhesive around the site (3M™ Vetbond™ Tissue Adhesive, 3M, Maplewood, Minnesota, USA). GI myoelectric activity was measured using up to six multi-contact planar electrodes (200 μm diameter, Micro-Leads Inc.; Fig. 1A). In six animals, the ventral abdominal vagal trunk was dissected free from the esophagus just below the diaphragm, and a flexible multi-contact cuff electrode (600-800 µm diameter; Micro-Leads, Inc) was placed around the trunk of the nerve. At time of implantation, the diameter of the nerve was measured intraoperatively and a nerve cuff of corresponding size was selected. Data from nerve stimulation are not included in this report. Four planar electrodes were secured to the ventral gastric surface using 8-0 silk suture. Additionally, in three acute preparations (Table 1: 29-18, 32-18 and 34-18) two additional planar electrodes were secured to the duodenum, placed at approximately 1.5 cm and 4 cm caudal to the pyloric sphincter, locations E and F, respectively. Ventral surface images were used to determine electrode location by drawing a triangle on top of the fat pad of the lesser curvature of the stomach (Fig. 1B). Left and right sides of the stomach were further divided by drawing a line from the mid-point of each side of the triangle, extended at a 90° angle (see Fig. 1B).

**Fig. 1:**
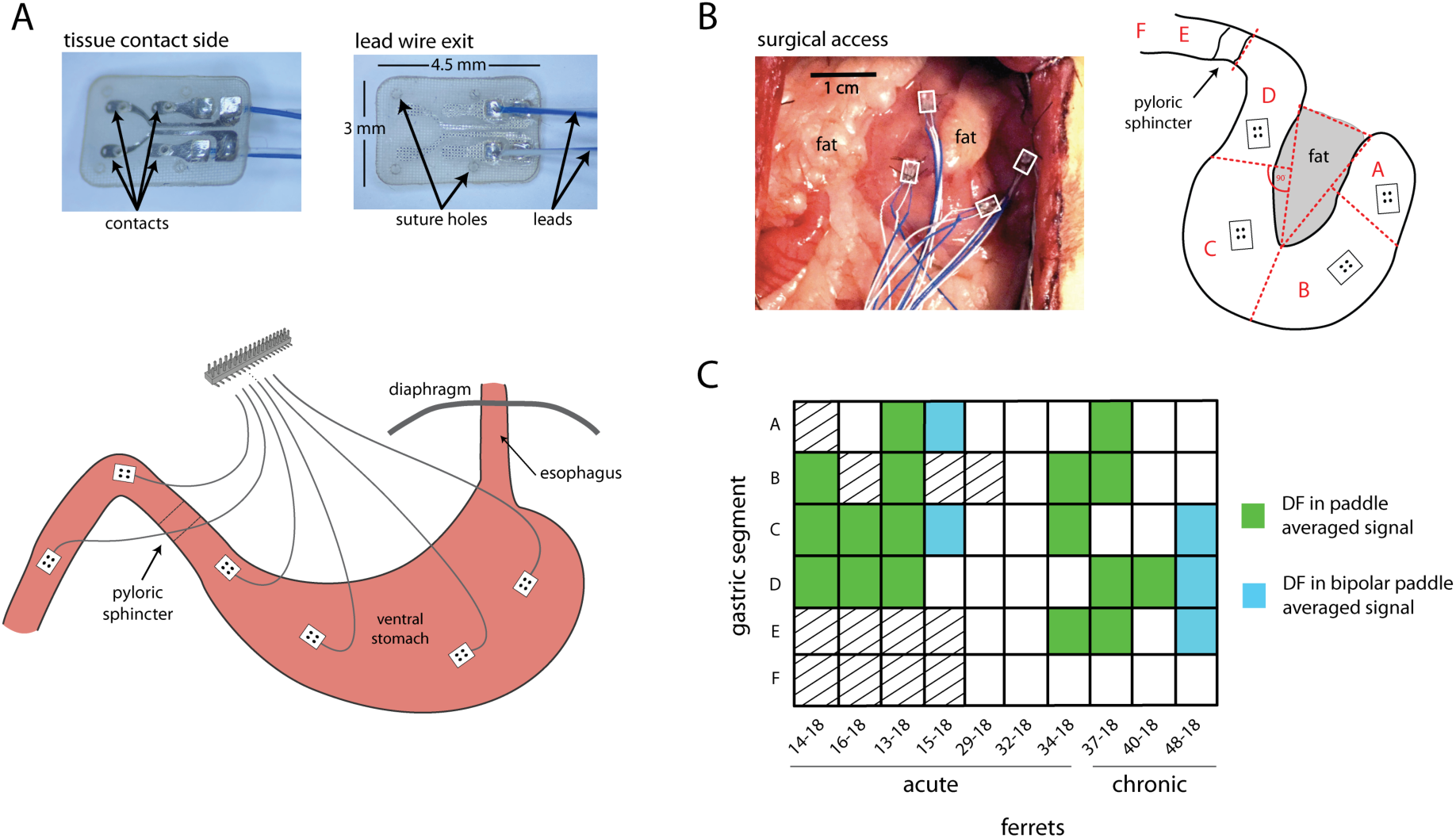
Placement of gastrointestinal recording electrodes. **A)** Micro-Leads planar electrodes, 1 to 6, were sutured to the stomach and duodenum of adult male ferrets. Planar electrodes are shown in inset images, which contain four contacts. **B)** A representative surgical placement of gastric electrodes 1 to 4 (top right), and the diagram shows how electrode position was determined (results in Table 1). **C)** GI myoelectric signals displaying dominant frequency are highlighted for paddle averaged signals (green) and bipolar differenced (blue).

### Chronic surgeries

Ferrets underwent a recovery surgery using aseptic techniques in a dedicated operating suite. Anesthesia was induced using an intramuscular injection of ketamine (15 mg/kg), and the animals were endotracheally intubated with either a 3.0 or 3.5 cuffed or uncuffed endotracheal tube. During surgery, anesthesia was maintained using isoflurane (1-2%) vaporized in O_2._Subcutaneous injections of sterile saline were used to replace fluid loss. A heating pad and infrared heat lamp were used to maintain rectal temperature (36–40°C). Animals were initially placed in the supine position, and the abdominal skin was sterilized using chlorohexidine and 70% isopropyl alcohol. After the animal was draped, the abdominal skin was sprayed with betadine solution.

Animals underwent a laparotomy to expose the abdominal contents. In a similar manner to that in the acute experiments, four 4-contact planar electrodes were placed on the stomach and two additional 4-contact planar electrodes were placed on the duodenum. Additionally, an infusion catheter was inserted into the stomach through a small incision on the lateral edge of the gastric fundus. The catheter was secured in place by tying a purse-string suture at the incision site and by placing Vetbond over the incision. The distal end of the gastric tube and the leads from the planar electrodes were routed subcutaneously, along the body flank and dorsal surface, to the neck using a trocar. The abdominal cavity was then lavaged with 60 ml of an antibiotic solution of Cefazolin diluted in sterile saline (1 gram: 100 cc saline; Cefazolin for injection, USP, Hi West-Ward Pharmaceuticals Corp, Eatontown, New Jersey, USA) to reduce the risk of infection. The abdominal musculature was sutured (2–0 silk; Ethicon), and the skin was closed with 3-0 monofilament (Ethicon).

The animals were rotated into the prone position and placed into a stereotaxic frame to secure the head. A midline, vertical, 6-cm incision was made on the skull. The skull was cleared of overlying musculature, and 4 to 8 self-tapping bone screws were inserted about the midline into the skull. Palacos bone cement (Zimmer, Warsaw, Indiana, USA) was placed over the bone screws and the electrode connectors were embedded in bone cement. The gastric tube was secured to the neck musculature using dacron and 4-0 silk suture. Post-surgical analgesia was provided for 72 h using buprenorphine (0.05 mg/kg, intramuscular). Amoxicillin (20 mg/kg BID) was administered orally for ten days after the surgery. Animals were weighed daily to assess body weight changes and were allowed to recover for at least 14 days before behavioral testing.

### Planar electrodes

Custom 4-contact paddle electrodes (Micro-Leads Inc.) were designed to conform to the stomach using a flexible silicone and platinum iridium 90/10 metal using a fusion-electrode substrate. The electrode contacts were 250 µm in diameter with pre-surgical impedances of 5-10 kΩ at 1 kHz. Suture holes were created through the silicone and nano-fiber reinforcement layers to prevent sutures from tearing the electrodes during chronic implantation (Fig.1A).

### Data acquisition

GI myoelectric signals were recorded from planar electrodes using a Grapevine Neural Interface Processor and a Nano2 recording headstage (Ripple LLC, Salt Lake City, Utah). Digitization of signals was performed directly on the headstage at 30 kHz with an input range of ±12 mV, resolution of 0.25 µV and a 0.1 Hz (6 counts per min) high-pass filter. Prior to all data collection, electrode impedances were recorded at 1 kHz using the Nano2 recording headstage. Additionally, in all acute preparations the intratracheal airway pressure (sampling rate (Fs)= 100Hz), EKG (Fs= 1kHz), blood pressure (Fs = 100Hz), gastric pressure (Fs = 100Hz), and rectal temperature (Fs = 100Hz) were sampled throughout the duration of the experiment using a CED Power 1401 16-bit analog to digital converter linked to a PC running Spike 2 version 7.15 software (Cambridge Electrode Design, Cambridge, UK).

In the acute preparation, baseline GI myoelectric activity was recorded at the onset of the experiment for up to 30 min. In 6 ferrets, mechanical distension of the stomach was achieved by infusing saline via the balloon catheter. The rate of infusion was set at 10 ml/min and the duration was varied to obtain 5, 10 and 20 ml of gastric distension across successive trials (GeniePlus Infusion Pump, Kent Scientific, Torrington, Connecticut, USA). For each trial, the stomach was held in the distended state for 2-5 min before the saline was drained at 10 ml/min. Additionally, in 6 ferrets, a bolus infusion of 5 mg/kg of intra-gastric emetine (emetine dihydrochloride hydrate, Sigma-Aldrich, St. Louis, Missouri, USA) was delivered and GI myoelectric activity was recorded for up to 60 min post-infusion to observe changes in GI myoelectric activity preceding retching and emesis.

For the chronic study, GI myoelectric activity was recorded using a cable tethered to the head connector in 3 freely moving ferrets during baseline control, intragastric infusion of water, and emetine. All ferrets were fasted 3 h prior to a recording session. Baseline GI myoelectric activity was recorded for up to 1 h for the first testing session, during which no food was provided. All animals were subsequently presented with food (Ensure Original Vanilla Flavor nutritional shake, Abbott Laboratories, Lake Bluff, Illinois, USA) for 30 min for at least 3 test sessions followed by at least 2 sessions in which vagal stimulation was applied while food was available for 30 min. Each ferret then underwent a trial with an emetic challenge in the form of 30 ml (5 mg/kg) of intragastric emetine and a control stomach distension trial in which 30 ml of water was infused. GI myoelectric activity was recorded for 1 h after infusion for both trials. Finally, animals 40-18 and 48-18 were subjected to a test session in which only vagal stimulation was applied for 30 min.

Chronic and acute recordings were analyzed post-hoc using MATLAB (Mathworks, Natick, MA) and Python (Python Software Foundation, https://www.python.org/) software. For every planar electrode, the waveform recorded on each of the 4 contacts was averaged to generate a single GI myoelectric waveform for that placement. Analysis of GI myoelectric activity was adopted from prior studies in awake behaving ferrets [14]. Briefly, each planar-averaged GI myoelectric signal was filtered using a low-pass Butterworth filter with a 2.5 Hz (150 cpm, 4^th^ order) cutoff. The filtered signal was then downsampled to 10 Hz and a second low-pass Butterworth filter with a cut-off frequency of 0.3 Hz (18 cpm, 2^nd^ order) was applied. In one ferret (13-18) non-physiological high amplitude transients that lasted 10-20 seconds were observed across all channels. These artifacts were removed by blanking the 1-min window around the artifact, prior to filtering and down-sampling. Each GI myoelectric signal was partitioned into 1-min segments and the power spectrum for each segment was obtained by computing the fast Fourier transform (FFT, bin size: 0.3 cpm). Each segment was characterized in terms of the dominant frequency (DF, frequency bin with the highest power in the 0 to 15 cpm range), total power in the 6-15 cpm range, and the percentage of total power in the bradygastric (from 1 to 3-cpm below DF), normogastric (between 1 cpm above and below DF) and tachygastric (from 1 to 3 cpm above DF) frequency bands [14]. To determine the DF of each GI myoelectric signal, a one-way ANOVA was conducted to determine if power in any frequency bins were statistically larger than others. Average power was calculated over all 1-min segments for individual frequency bins. A post-hoc multiple comparison was then performed for pairs of global average power peak and each of the local peaks simultaneously, for testing the existence of a statistically significant DF peak. The detection of DF was done in R (version 3.5.1, R Foundation for Statistical Computing, Vienna, Austria). For emetine infusion trials, the dominant frequency and percentage of power in the normogastric range (P_norm_) prior to emetine infusion was compared to that after emetine infusion up to the first retch. Similarly, for balloon distension trials, pre-distension DF and P_norm_ were compared to that during the hold phase of distension.

### Machine learning

For each GI myoelectric signal, windowed features were obtained for 1-min segments described previously. Per segment, the power in the brady (P_brady_), normo (P_norm_) and tachygastric (P_tachy_) ranges, dominant frequency (DF) and power within the 0.3 cpm band for the dominant frequency (DP) were extracted along with line length (LL, sum of the magnitude of the first signal derivative over time) and zero crossing features (ZX, number of times the algebraic sign of the signal changed). All features were normalized to the median value per window. Because multiple features rely on the presence of a DF in the GI myoelectric signal, animals were excluded from machine learning analysis if they did not exhibit a DF for any paddle averaged signal or any bipolar pair of paddle averaged signals. For animals that exhibited a DF, the performance of a support vector machine (SVM), k-nearest neighbor (kNN) and naïve Bayes classifier trained independently for each subject was compared for detecting gastric state during the emetine trial. Prior to training, parameters for each algorithm (number of neighbors for kNN and kernel, gamma and C for SVN) were determined via a grid search implemented using the *Scikit*-*learn* library in Python (https://scikit-learn.org). The GI myoelectric activity was partitioned into a pre-infusion ‘baseline’ state, a post-infusion ‘early’ state, and a pre-retch ‘late’ state. GI myoelectric activity recorded prior to emetine infusion was labeled baseline. GI myoelectric activity recorded in the interval between emetine infusion and the first retch was partitioned in half into two equal intervals of time (i.e. ‘early’ and ‘late’). 5-7 min of baseline GI myoelectric activity was collected for 3 ferrets (13-18, 16-18, 15-18) and less than 1 min of baseline was collected for the remaining ferret (14-18). In these ferrets, training and testing was performed on two states, excluding baseline. One additional ferret (32-18) was excluded from classification analyses because an electric heating pad induced excessive noise in the recorded signals. During classifier evaluation, to avoid class imbalances, the number of time windows used per gastric state were kept equal. This meant the number of time windows per class was limited by the shortest recorded gastric state.. Feature sets were selected using a greedy stepwise process {Hocking, 1976 #3697}. At each step, the classifier was trained on one additional feature and a 5-fold cross validation was carried out. Features with the highest cross-validation accuracy were retained at each step. Additionally, the chance level of prediction for each animal was established by repeating 5-fold cross-validation after randomly scrambling class labels.

## Results

### A: Acute anesthetized ferrets

#### Baseline GI myoelectric activity

Across the 7 acute preparations a total of 34 GI myoelectric paddles were placed on the serosal surface of the stomach. Fourteen paddles (Fig. 1D) corresponding to animals 14-18, 16-18, 13-18, and 34-18 displayed a statistically significant (p<0.0001) dominant frequency peak at 9.53 ± 0.67 cpm. For the remainder of the animals the paddle averaged GI myoelectric signal from each paddle did not display a DF. However, in animal 15-18 the bipolar difference of the averaged GI myoelectric signal was calculated for all possible bipolar pairs and 2 out of the 6 possible bipolar pairs reported a DF of 8.55 ± 0.15 cpm. These paddles were located in gastric segments A and C. For animals 14-18, 16-18, 13-18 and 34-18, the DF was invariant to the location of the GI myoelectric signal. The DF observed for segments A, B, C, D and E was 9.75 ± 0.64, 9.60 ± 0.95, 9.53 ± 0.67, 9.70 ± 0.35 and 8.4 cpm across animals. For all subsequent analysis, data from these 5 animals were used. For the same 14 paddles that showed a DF, the P_norm_ at baseline was 57.1 ± 14.4 % across all locations. This translated to 50.4 ± 22.8%, 60.9 ± 6%, 59.0 ± 11.6%, 50.5 ± 6 % and 75.7% P_norm_ for segments A, B, C, D and E. Figure 2 shows an example of baseline recording for animal 13-18 with the power spectrum of the gastric myoelectric activity recorded at segment C.

**Fig. 2:**
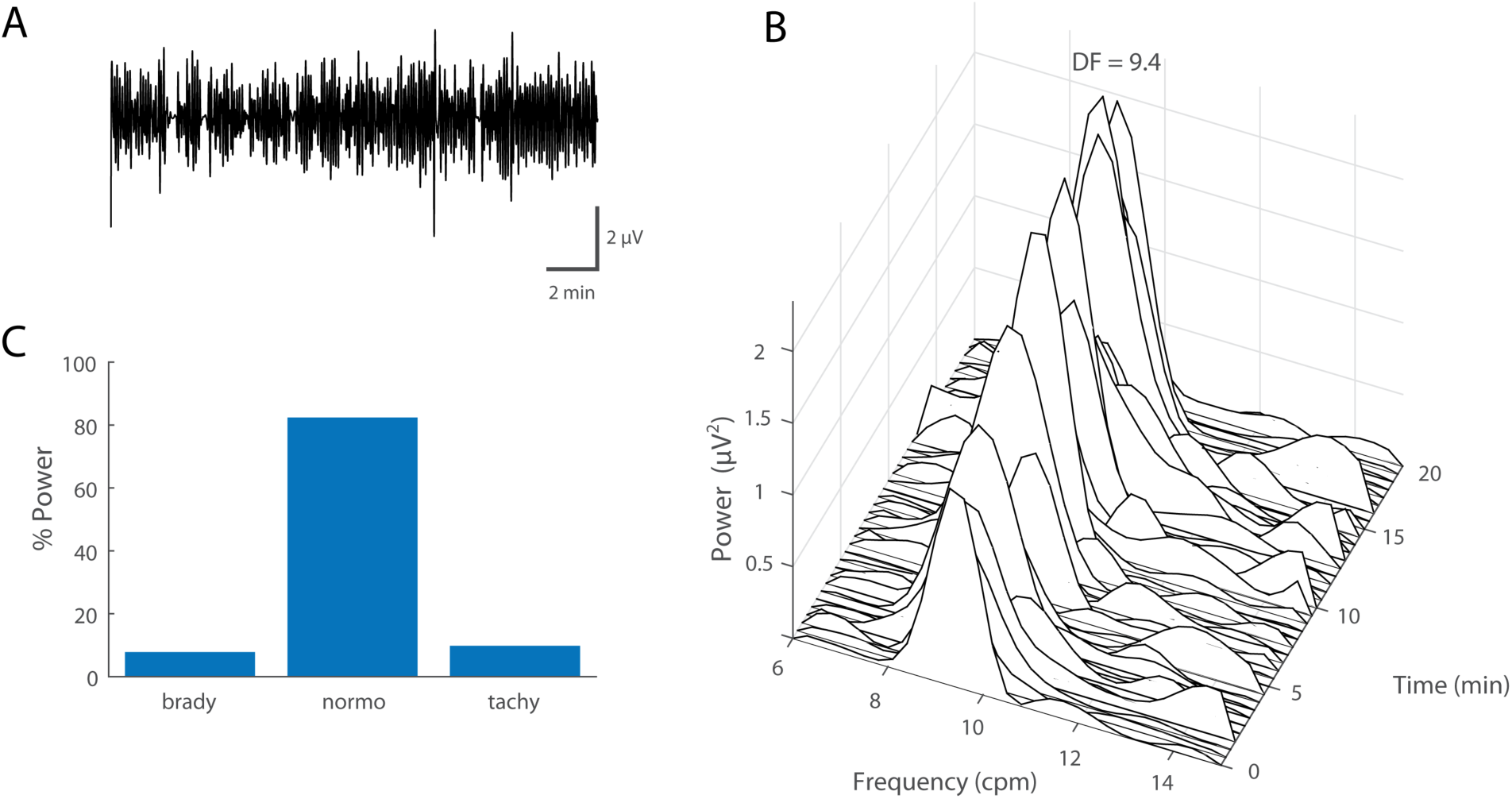
Baseline electrogastrogram (GI myoelectric) in anesthetized ferret. **A)** Example of a 20 min filtered and downsampled GI myoelectric recorded from segment 1 of an anesthetized ferret at baseline. **B)** Waterfall plot of the power spectral density for the waveform shown in A. Each time window corresponds to the FFT of 1 min of GI myoelectric data. **C)** Percentage of total power in the 6-15 cpm range partitioned by bradygastric (6.4 − 8.4 cpm), normogastric (8.4 − 10.4 cpm) and tachygastric (10.4 − 12.4 cpm) ranges for a signal with a DF of 9.4 cpm at baseline.

#### Effect of gastric distension on GI myoelectric activity

Distension trials were carried out in 4 of the 5 ferrets (14-18, 15-18, 16-18, 34-18). Figure 3 shows an example of GI myoelectric activity recorded during gastric distension at 20 ml. Across ferrets, changes in GI myoelectric activity during the distension phase of gastric distension were compared to baseline GI myoelectric activity collected immediately prior to distension (Fig. 3C).

**Fig. 3:**
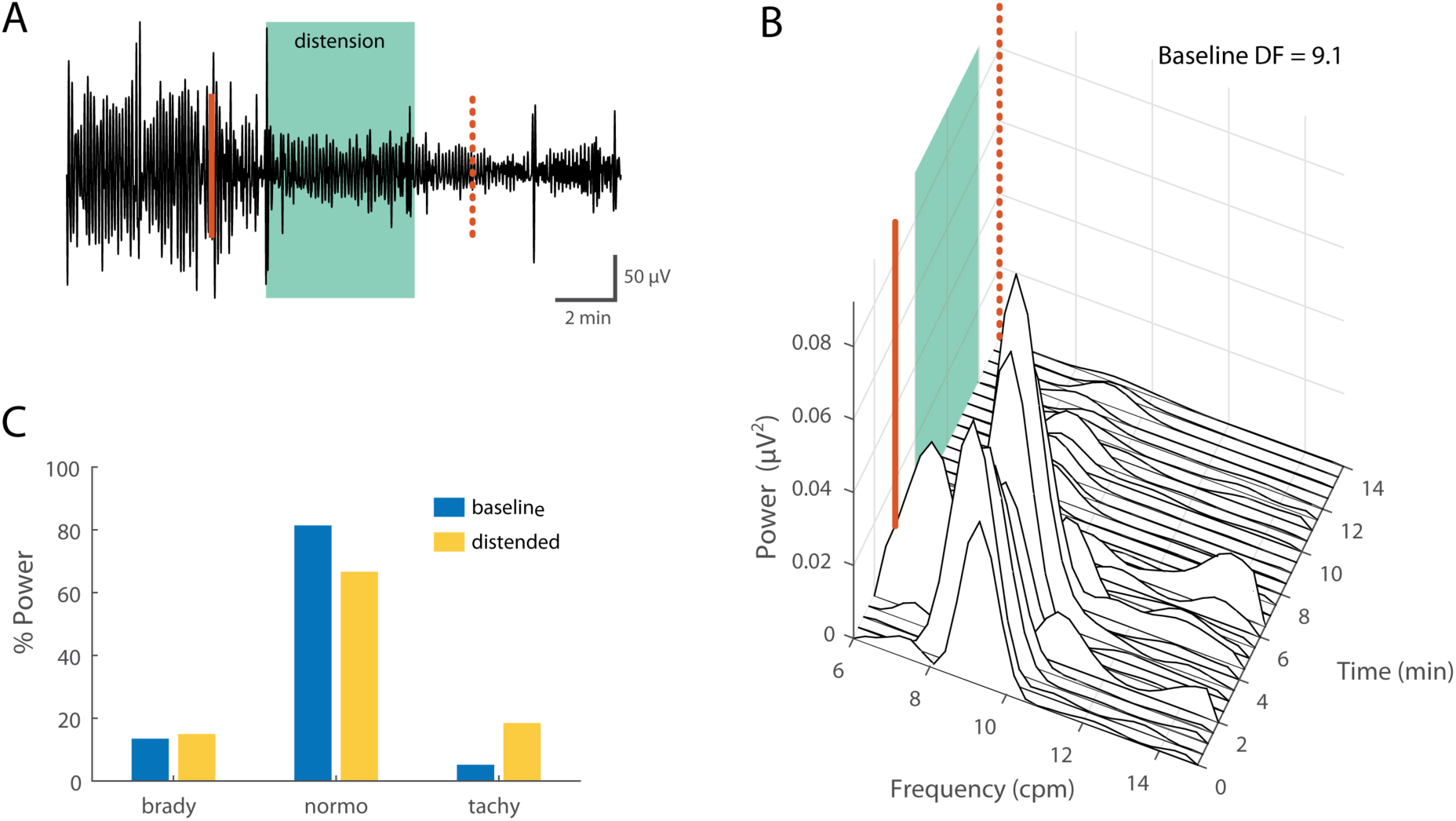
Effect of gastric distension on GI myoelectric. **A)** Example of a 19 min filtered and downsampled GI myoelectric recorded from segment 3 of an anesthetized ferret at during gastric distension at 20 ml. Distension is maintained for 5 min (green shaded area) between infusion start and end (solid and dashed line). **B)** Waterfall plot of the power spectral density for the waveform shown in A. Each time window corresponds to the FFT of 1 min of GI myoelectric data. **C)** Percentage of total power in the 6-15 cpm range partitioned by bradygastric (6.1 − 8.1 cpm), normogastric (8.1 − 10.1 cpm) and tachygastric (10.1 − 12.1 cpm) ranges for a signal with a DF of 9.1 cpm at baseline versus during distension.

For low volume gastric distension (5 ml) ferret 14-18 displayed a 10-20% decrease in the DF from baseline across all gastric segments (Fig. 4A). Animal 16-18 displayed no change in the DF at segment C, a 10% increase in the DF at segment D and a 10% decrease in the DF at segment A. ferret 34-18 showed a 50% increase in the DF at the duodenum whereas only a 10% change in the DF at segment C and 4. At 10 ml distension, animals 16-18, 15-18 and 34-18 displayed a 10-50% decrease in DF across all gastric segments. Animal 14-18 in this instance showed an opposite trend where the DF at segments B and D showed an increase in DF. At the maximum volume of distension animals 14-18 and 16-18 both displayed up to a 40% increase in the DF across all gastric segments. Interestingly, the trends in DF were not mirrored in the P_norm_ (Fig. 4B). For distension at 5 ml, all gastric segments in animal 14-18B showed a 50% decrease. Segment D in animal 16-18 also showed a 50% decrease in P_norm_ while segment A showed a 10% increase in the P_norm_. ferret 34-18 showed a 50% increase in P_norm_ at the segment E while segments B and C showed a 50% increase in P_norm_. Distension at 10 ml, did not show any uniform trend across subjects or locations, however 20 ml distension seemed to have the opposite effect on P_norm_ as 5 ml distension for animals 14-18 and 16-18. In 2 animals (29-18 and 32-18), distension at 20 ml induced retching and emesis however since no DF was observed on any GI myoelectric recording at baseline these animals were excluded from subsequent analyses.

**Fig. 4:**
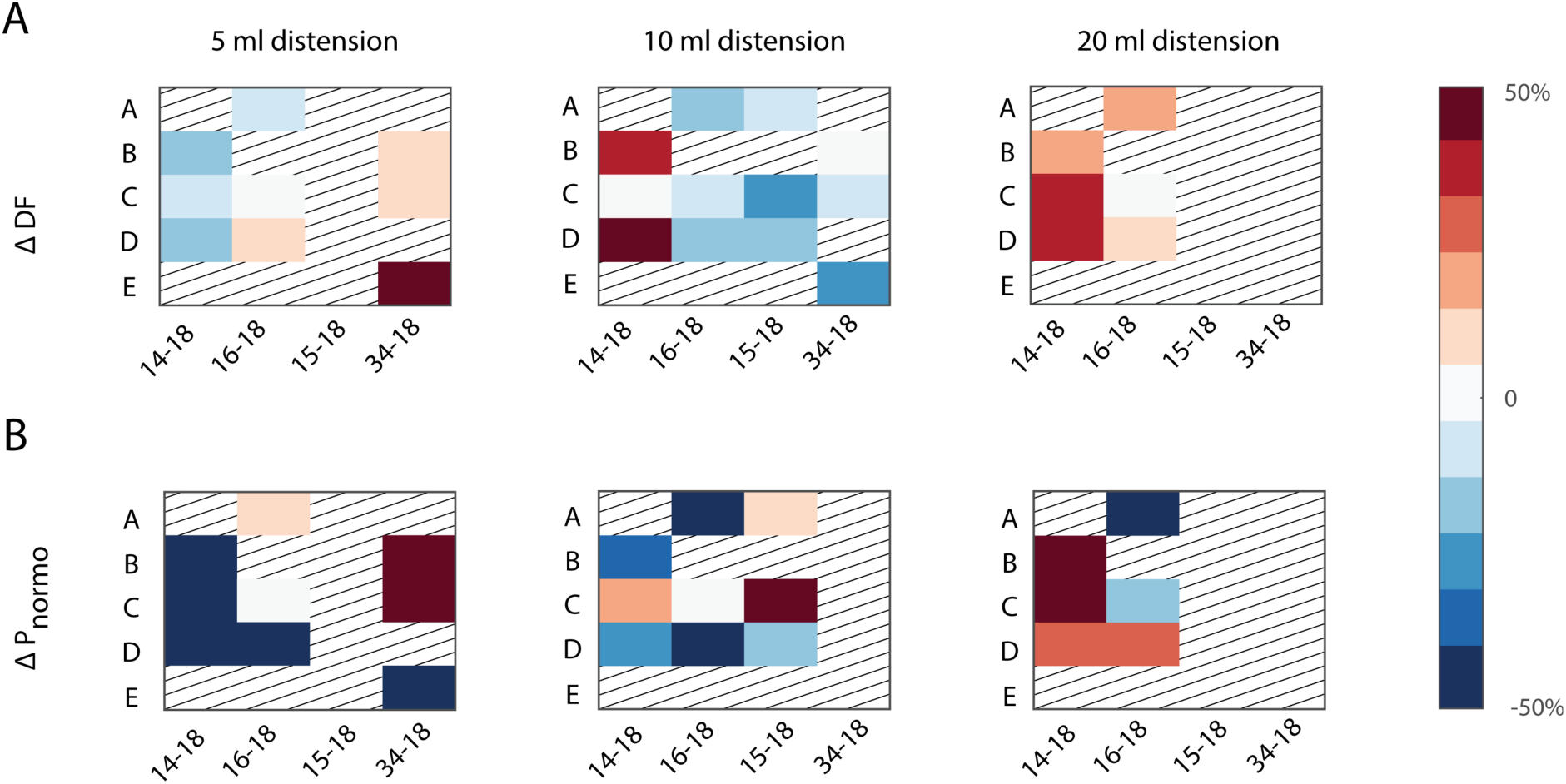
Effect of gastric volume on DF and P_norm_. **A)** Heatmap of the change in DF and **B)** percentage of total power in the normogastric range in response to gastric distension at 5, 10 and 20 ml (columns) across subjects and gastric segments for animals B, C, D, E, H. Grating represents gastric segments that did not display a DF at baseline or trials that were not administered.

#### Effect of emetic stimuli on GI myoelectric activity

Emetine infusion was carried out in 4 of the 5 ferrets (14-18, 16-18, 13-18, 15-18) Across these ferrets, emetine induced retching within 29.5 ± 2.9 min of infusion. Following emetine infusion there was a gradual decrease in P_norm_ across most gastric segments that persisted until the peri-retch (Fig. 5A). Only segment D in animal 13-18 showed a 50% increase in the P_norm_ in the lead up to retching. For all ferrets, gastric segments B and C, showed up to a 30% increase in the DF (Fig. 5B) whereas segment A showed a less than 10% decrease in the DF. Segment D appeared show more variability since DF increased in animal 16-18 and 15-18, decreased in 14-18 and remain unchanged in 13-18. Figure 6 shows an example of GI myoelectric activity recorded from segment 3 during emetine infusion.

**Fig. 5:**
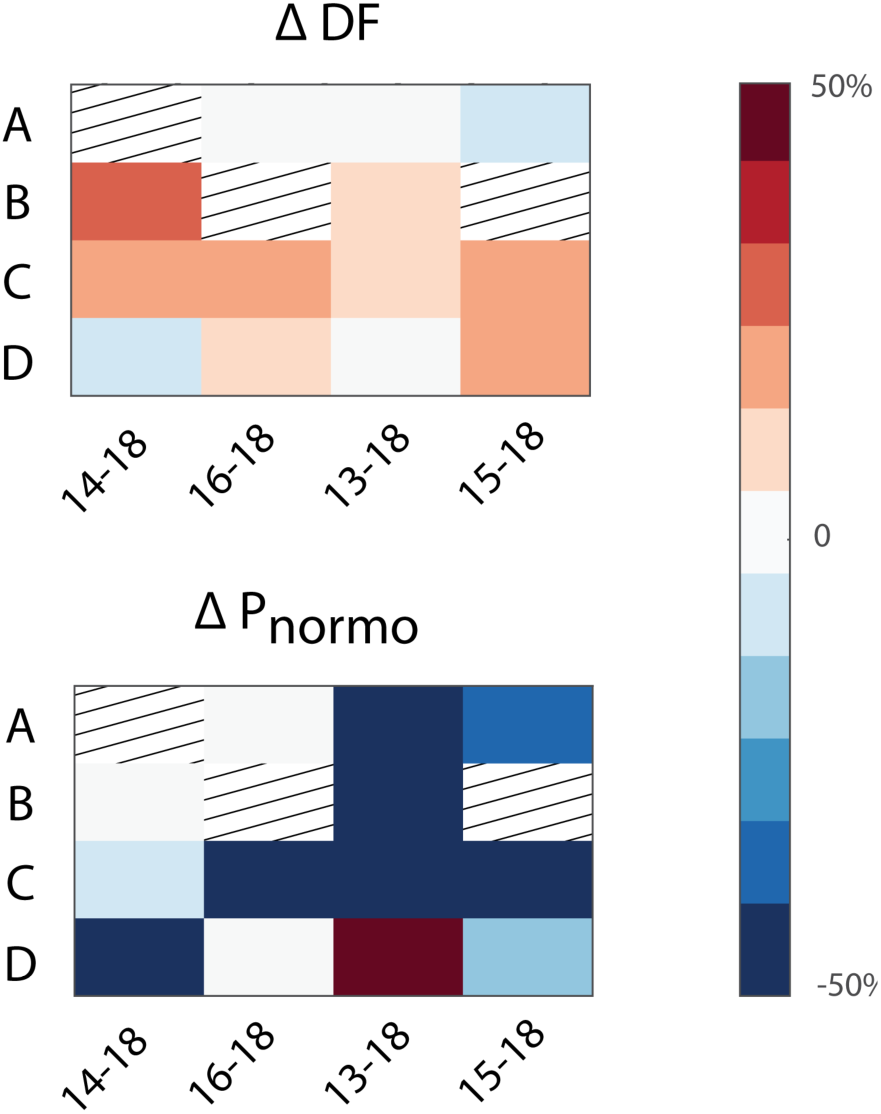
Change in DF and P_norm_ after intra-gastric emetine. **A)** Heatmap of the change in DF and **B)** percentage of total power in the normogastric range after emetine infusion across subjects and gastric segments for animals 14-18, 16-18, 13-18 and 15-18. Grating represents gastric segments that did not display a DF at baseline or trials that were not administered.

**Fig. 6:**
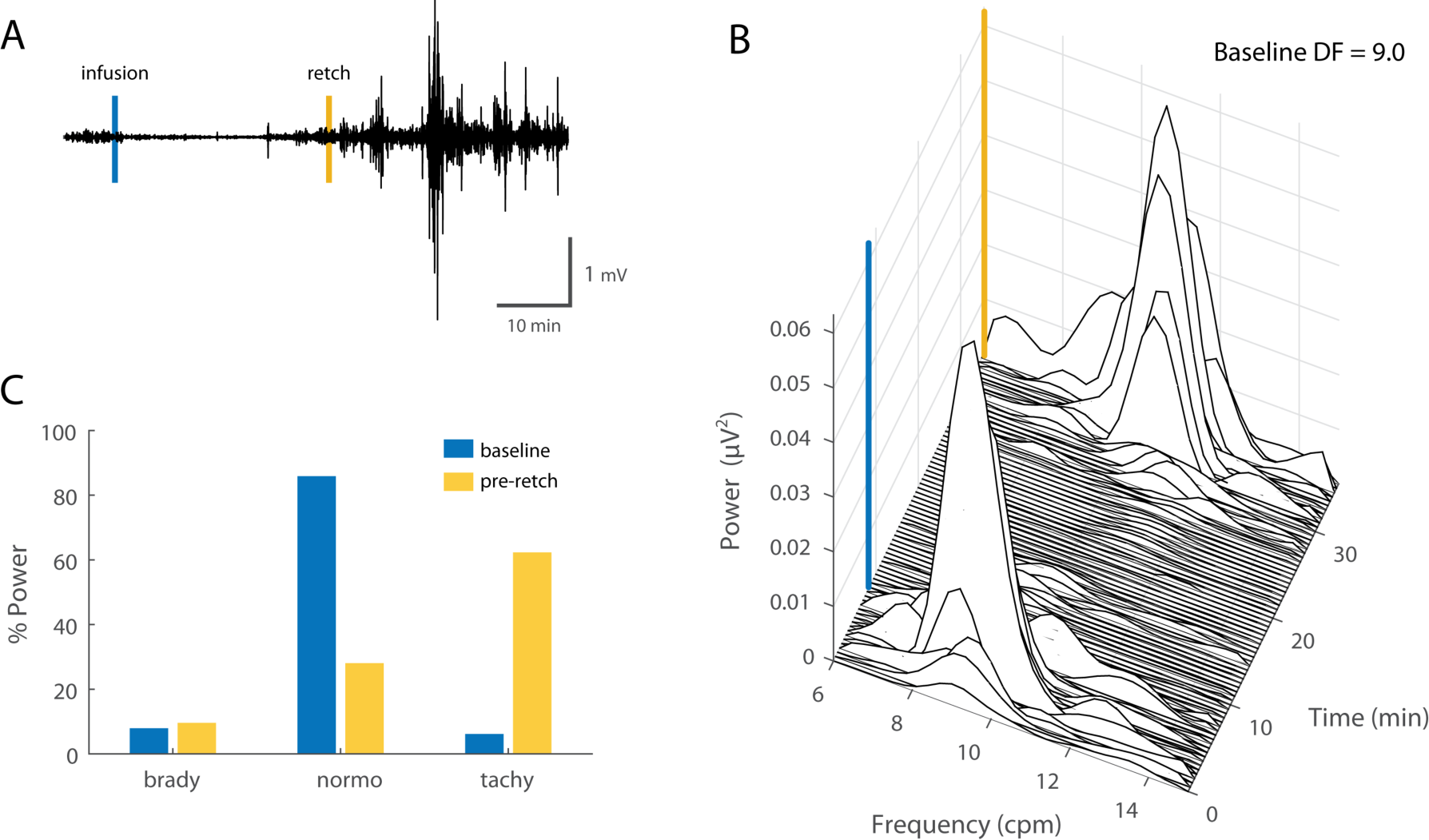
GI myoelectric activity recorded from segment 3 during emetine infusion. **A)** Example of a 70 min filtered and downsampled GI myoelectric recorded from segment 3 of an anesthetized ferret during emetine infusion trials. Emetine induced retch (yellow line) is observed approximately 28 min post emetine infusion (blue line). **B)** Waterfall plot of the power spectral density for the waveform shown in A. Each time window corresponds to the FFT of 1 min of GI myoelectric data. **C)** Percentage of total power in the 6-15 cpm range partitioned by bradygastric (6 − 8 cpm), normogastric (8 − 10 cpm) and tachygastric (10 − 12 cpm) ranges for a signal with a DF of 9 cpm at baseline versus 20 min prior to the first emetine induced retch.

### B: Chronically implanted ferrets

#### Baseline GI myoelectric activity

In the chronic preparations, of the 18 paddles that were implanted only 5 paddles across 2 animals showed a significant DF in the paddle averaged signal. GI myoelectric activity recorded from ferret 37-18 showed a DF peak at gastric segments A, B, D and E. The average DF observed across these paddles was 9.53 ± 0.13 cpm. In ferret 40-18, segment D showed a DF of 9.6 cpm at baseline. No other paddle showed a DF peak at baseline for this animal. However, in ferret 48-18 the bipolar difference of the averaged GI myoelectric signal was calculated for all possible bipolar pairs and 2 bipolar pairs reported a mean DF of 8.25 ± 1.05 cpm Throughout the duration of the 1-hour recording, the DF and the power spectrum of the GI myoelectric activity displayed variability across consecutive windows. The mean DF observed in awake behaving animals (9.54 ±0.12 cpm) was not significantly different from that observed during acute experiments (9.17 ± 0.88).

#### GI myoelectric activity during feeding

For feeding trials, baseline GI myoelectric activity was recorded for 10 min prior to food presentation. There was a high intra and inter subject variability in the rate and volume of consumption of food per trial. Nevertheless, food intake produced an immediate reduction in the total power in the 6-15 cpm range resulting in a near flat-lining of the GI myoelectric activity. The change in DF and P_norm_ across gastric segments showed variability and there were no common trends observed across subjects. For animal 37-18, GI myoelectric activity recorded during the first feeding trial displayed an increase in the DF across all gastric segments. This increase was highest for segment B at 30% of baseline. However, subsequent feeding trials showed no change or a slight decrease in DF after food consumption for all segments. Interestingly, changes in P_norm_ showed a more consistent trend across recording sessions. Segment A and B seemed to follow the opposite trend in terms of changes in P_norm_. For 2 of the 4 feeding trials the P_norm_ remained unchanged for both segments. For the remainder of the feeding trials segment B showed a 25% decrease in P_norm_ while segment A showed a 25% increase for the same trials. The overall suppression of power in the GI myoelectric activity persisted after the ferret had stopped food consumption and up to 10 min after the food was removed, suggesting that satiation and not just dilation of the stomach may have a role in the changing GI myoelectric activity observed in this animal.

For ferret 40-18, segment D was the only paddle averaged signal to display DF displayed variability in the direction of DF change during feeding and emetine trials. For early feeding trials, there was a 0-5% increase in the DF whereas subsequent feeding trials resulted in a 5-15% decrease in the DF. Interestingly, this trend was reflected in the P_norm_ for this animal. GI myoelectric activity from segment D showed a 50-125% increase in the P_norm_ during the first two feeding trials however subsequent feeding trials showed a decrease in the P_norm_. Ferret 48-18 also showed a similar variable trend in the DF as well as P_norm_ across feeding trials. Figure 7 shows an example of gastric myoelectric activity recorded during food consumption in an awake behaving ferret

**Fig. 7:**
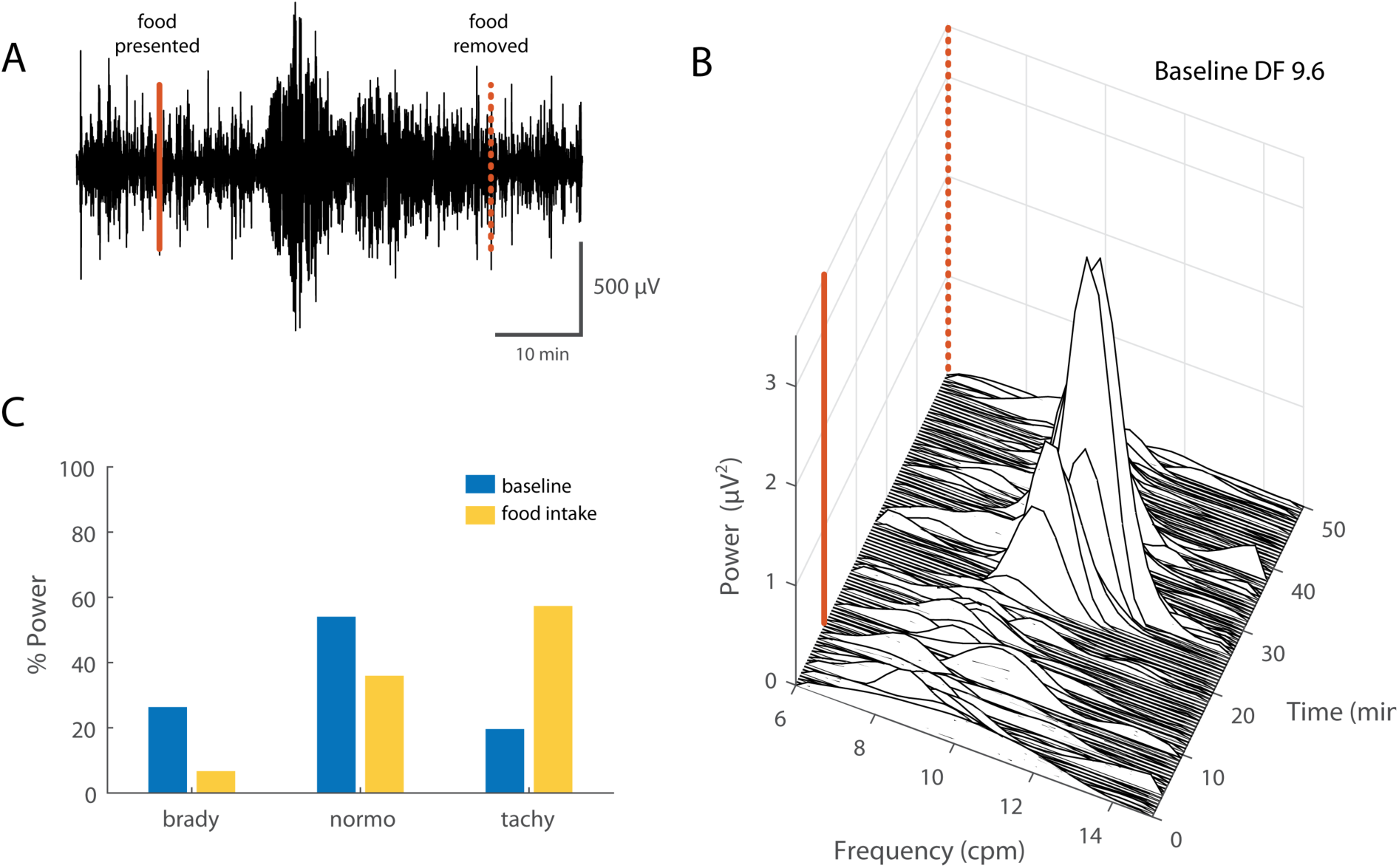
Gastric myoelectric activity during food consumption. **A)** Example of a 60 min filtered and downsampled GI myoelectric recorded from segment 4 of an awake behaving ferret during a feeding trial. Solid and dashed red lines denote when food was presented and withdrawn. **B)** Waterfall plot of the power spectral density for the waveform shown in A. Each time window corresponds to the FFT of 1 min of GI myoelectric data. **C)** Percentage of total power in the 6-15 cpm range partitioned by bradygastric (6.6 − 8.6 cpm), normogastric (8.6 − 10.6 cpm) and tachygastric (10.6 − 12.6 cpm) ranges for a signal with a DF of 9.6 cpm at baseline versus during food presentation and consumption.

#### Effect of emetic stimuli on GI myoelectric activity

For emetine trials, baseline GI myoelectric activity recorded prior to emetine infusion was compared to pre-retch GI myoelectric activity. Across all 3 chronic animals the mean interval between emetine infusion and the first retch was 23.7 ± 2.5 min. For 37-18, DF at gastric segments A, B, D and E showed a 5-10% decrease. In terms of P_norm_, there was no change at gastric segment B and E whereas segments A and D displayed a 50-75% increase in the P_norm._

For 40-18, two emetine trials were carried out on separate testing days, with one week between tests. For the first emetine trial segment D showed a 5-15% change in the DF from baseline. For the second emetine trial, segment 2 showed a 5% decrease in the DF while it remained unchanged during the second emetine trial. P_norm_ at segment D showed a 25% increase during the first emetine trial and showed a 25% decrease in P_norm_ during the second emetine trial. while segment B remained unchanged for both emetine trials.

For animal 48-18 the trend in DF was similar to that of P_norm_. Segment C and E showed no change in the DF while segment D showed a 5% decrease in the DF while the rest of the segments remained unchanged. For segment D the P_norm_ displayed a 25-50% decrease while the rest of the segments remained unchanged. Figure 8 shows an example of GI myoelectric activity recorded during emetine infusion in an awake behaving ferret.

**Fig. 8:**
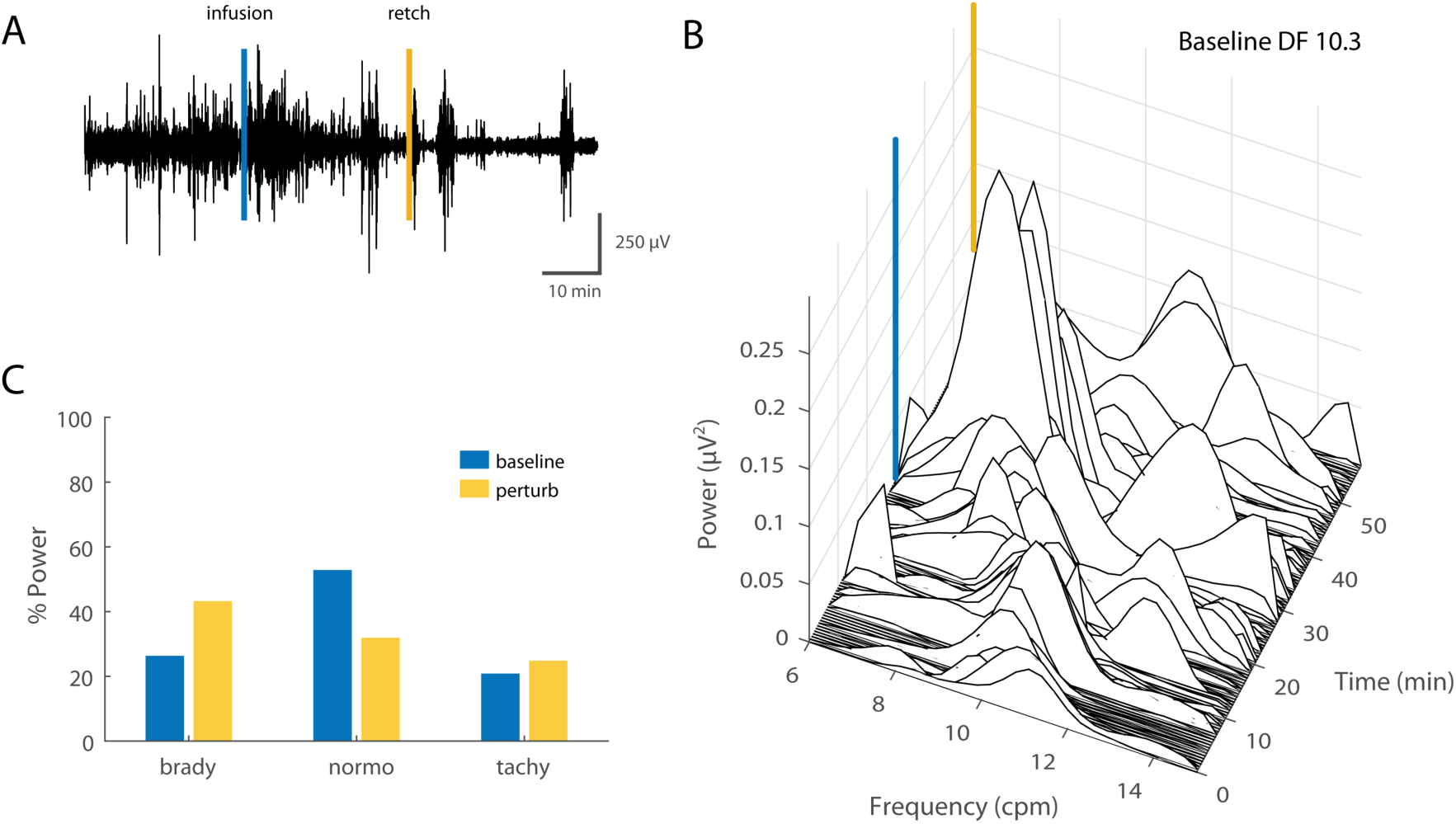
GI myoelectric activity recorded during emetine infusion in an awake behaving ferret. **A)** Example of a 90 min filtered and downsampled GI myoelectric recorded from segment 1 of an awake behaving ferret during an emetine infusion trial. Emetine induced retch (yellow line) was observed approximately 27 min post infusion (blue line). **B)** Waterfall plot of the power spectral density for the waveform shown in A leading up to the first retch. Each time window corresponds to the FFT of 1 min of GI myoelectric data. **C)** Percentage of total power in the 6-15 cpm range partitioned by bradygastric (7.3 − 9.3 cpm), normogastric (9.3 − 11.3 cpm) and tachygastric (11.3 − 13.3 cpm) ranges for a signal with a DF of 10.3 cpm at baseline versus pre-retch

#### Effect of gastric distension on GI myoelectric activity

Baseline GI myoelectric activity was recorded for up to 10 min prior to gastric distension in animals 40-18 and 48-18. Similar to previous trials, the effects were varied across subjects. For ferret 40-18, the DF at gastric segment D remained unchanged (Fig 9A). Similarly for animal 48-18 distension was accompanied by a 5-10% increase in the DF at segment C whereas segments D and E remained unchanged. For ferret 40-18 the P_norm_ gastric distension resulted in a 50% decrease in P_norm_ at segment D whereas the opposite effect was seen in ferret 48-18 where gastric distension resulted in a n increase in P_norm_ at segments C and E while the P_norm_ remained unchanged for the segment D (Fig. 9B).

**Fig. 9:**
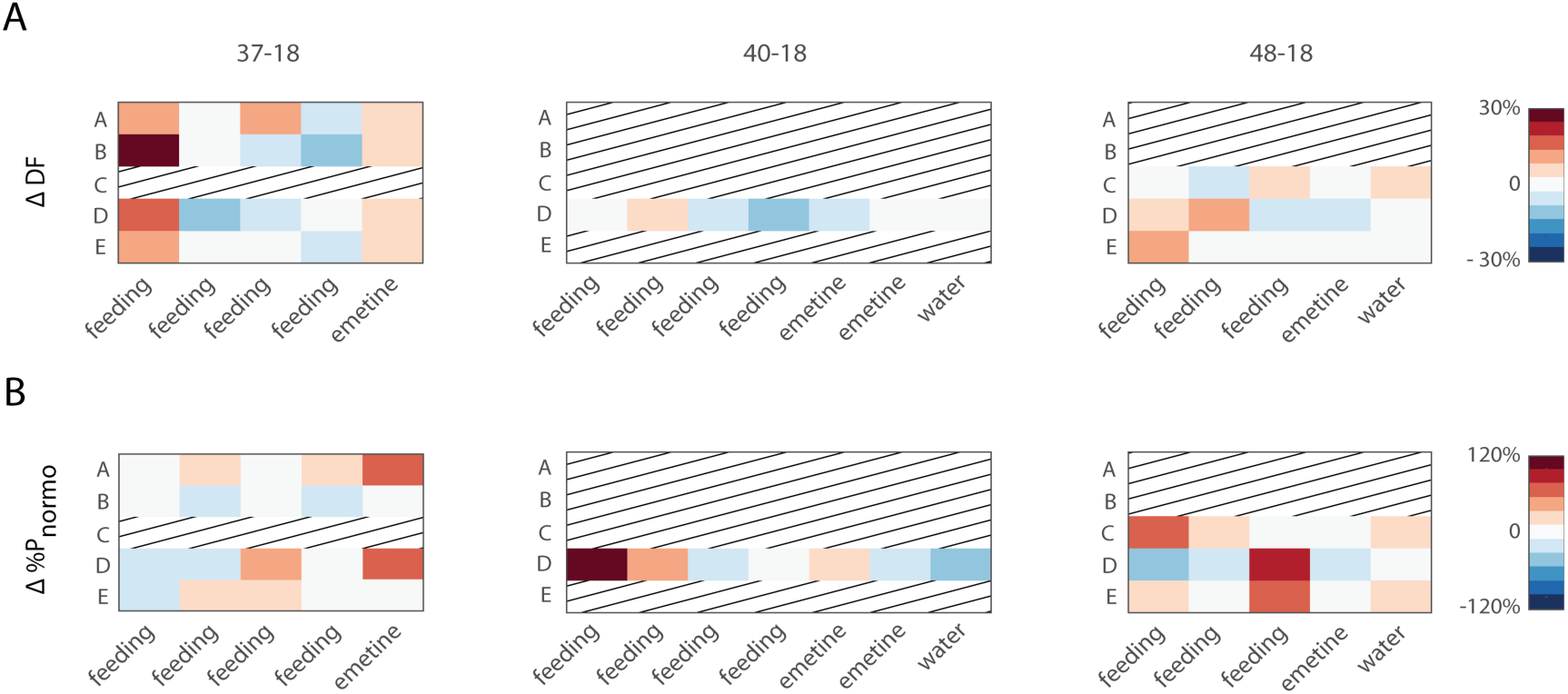
Change in DF and P_norm_ during gastric distension. A) Heatmap of the change in DF and B) percentage of total power in the normogastric range across multiple days of testing for subjects 37-18, 40-18 and 48-18 (columns).

### C: Detecting gastric state using standard machine learning algorithms

The performance of each learning algorithm was evaluated per subject and the greedy stepwise process was used to identify feature subsets that produced the highest classification accuracy (Table 2). For ferrets 14-18 no baseline data were collected prior to emetine trials therefore classification was only between early and late pre-retch. For this ferret, the overall testing accuracy was above chance (82%). For animals 16-18, 13-18, and 15-18 the classifiers were trained on 3 gastric states therefore chance level for prediction was 33% while testing accuracy for all 3 animals was above 80%. Similar to data from acute anesthetized experiments there was substantial inter-subject variability in terms of the optimal features and GI myoelectric activity signals required to detect gastric state. Across subjects, optimal features typically included one or more bandpower feature. The greedy algorithm also demonstrated that inclusion of time domain features (LL and ZX) during training, greatly improved classifier performance across all subjects. Interestingly, although there were no uniform trends in change in DF and P_norm_ following emetine infusion, the gastric segment with the highest predictive power was segment C across animals 14-18, 16-18, and 13-18. For animal 15-18 the bipolar paddle differenced GI myoelectric activity recorded from segment A displayed the highest predictive power.

**Table 2:** Optimal classifier performance.

| subject | accuracy | algorithm | features | segments |
| --- | --- | --- | --- | --- |
| **14-18** | 82% | kNN | $P_{\text{norm}}$, $P_{\text{tachy}}$,<br>DF, ZX | C |
| **16-18** | 85% | SVM | $P_{\text{tachy}}$, LL, ZX | C |
| **13-18** | 82% | kNN | $P_{\text{brady}}$, $P_{\text{norm}}$,<br>LL | C |
| **15-18** | 81% | SVM | LL, ZX | A-A |

A confusion matrix was constructed for classification accuracy collapsed across all subjects. For subjects 16-18, 13-18, and 15-18 (fig 10A, 3 gastric states) and 14-18 (2 gastric states). Consistent with individual subject results, the main diagonal of the confusion matrix for subjects 16-18, 13-18 and 15-18 showed an approximately 80% classification accuracy across subjects. The ‘early’ state labels had the lowest classification accuracy and was frequently mislabeled as a ‘late’ state. For the 2-state confusion matrix, the average classification accuracy was approximately 82% and ‘early’ state was mislabeled as ‘late’ state more often than the reverse.

**Fig. 10:**
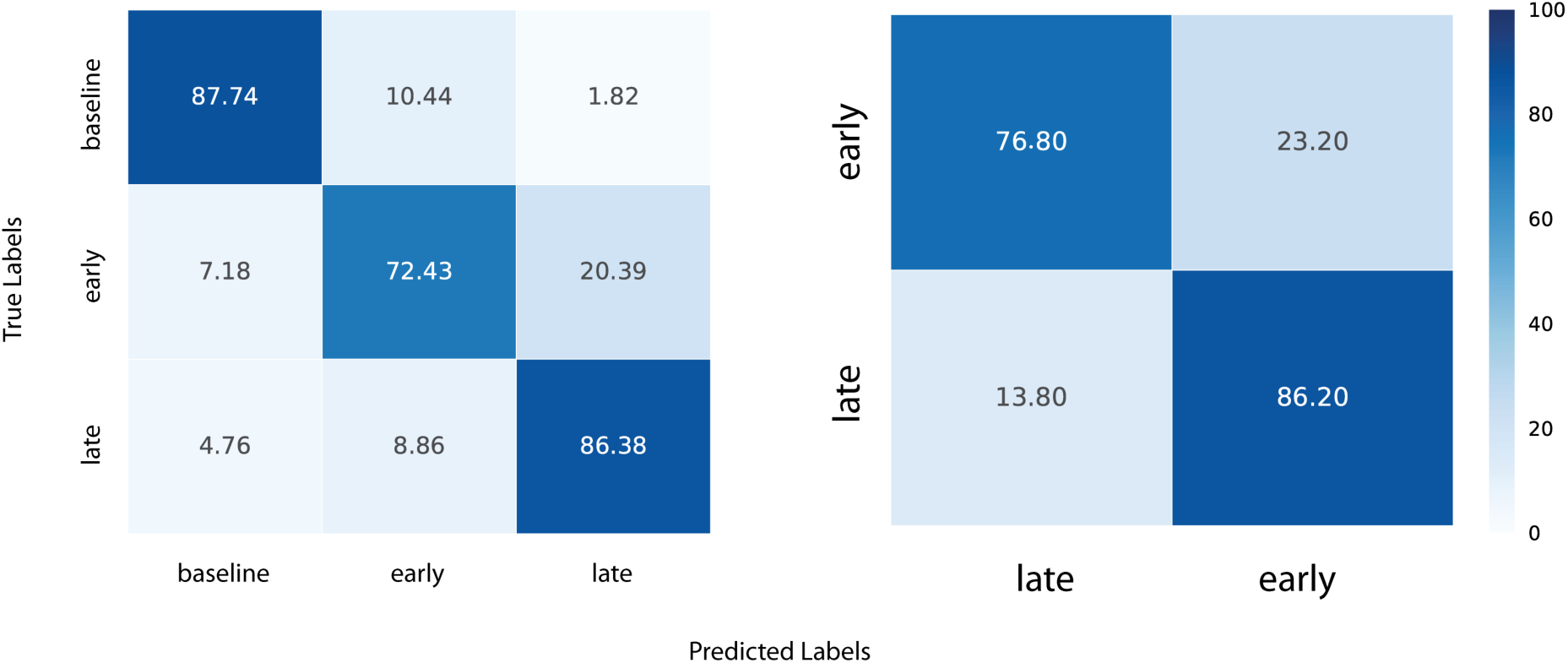
Confusion matrix. Aggregated confusion matrix for optimal feature set and best performing classifier across all subjects 3-state (13-18, 15-18 and 16-18) and 2-state (14-18) classification

## Discussion

The present study is the first demonstration of machine learning algorithms used to detect the physiological state of the stomach and onset of retching in ferrets. In the acute and chronic experiments, the existence of a DF was used as a criterion to include or reject recorded GI myoelectric signals. This criterion was based on prior studies involving awake behaving ferrets, dogs and mice [14, 19–21] that described the DF as a characteristic and consistent peak in the power spectrum of myoelectric signals recorded across the stomach. Five animals in the acute study displayed a DF in the GI myoelectric signals for at least one gastric segment and the intra-subject variability of the DF (across paddles) was low. Interestingly, gastric segments C and D (Fig. 1C) in the present study showed a DF peak across multiple ferrets. It is worth noting that prior work in ferrets [14] has focused on GI myoelectric signals recorded from the stomach wall in regions that can be roughly aligned to gastric segment C or D in the present study.

For emetine infusion and gastric distension of the stomach, there was no common trend across animals in terms of the shift in DF or the change in P_norm_. It is possible that, single trials of emetine or gastric distension led to long term or even permanent (for the duration of the experiment) changes in GI myoelectric physiology that obscured any possible trends. In animals 14-18, 16-18, 13-18, and 15-18, electrical stimulation of the vagus nerve was performed to measure the effects on GI myoelectric signals. Additionally, microelectrode arrays were implanted in the nodose ganglion to monitor single unit activity in response to gastric perturbation. We chose not to report these effects here because these manipulations were in a smaller subset of animals or were not amenable to machine learning. However, it is possible that these procedures also disrupted normal afferent signaling and GI myoelectric responses during the experiment. Furthermore, emetine infusion, vagus nerve electrical stimulation, and gastric distension were not administered in the same sequence across all acute preparations (Fig. 11). This variability in experiment design itself may be a confound that explains the observed variability in GI myoelectric activity across animals. In the future, repeated trials of the same perturbation will have to be applied per animal to identify whether the observed signal variability is truly stochastic or is a consequence of the gastric perturbation. Unlike prior work in awake behaving animals this study found that incidence of a DF in the GI myoelectric signal is highly variable. For the acute preparations this variability may be due to administration of anesthesia, however the absence of DF in several chronic recordings indicates that DF may not always be a reliable biomarker of GI state. Inconsistency in DF may also be because our placement of electrodes differed in some animals (Table 1). This was due to the difficultly in observing landmarks that would guide electrode placement during acute surgery, although planar electrode placement was largely similar across many of the animals. Future work will have to focus on developing a biomarker of GI myoelectric signals that can be reliably detected and is clearly modulated when the stomach is perturbed via mechanical, chemical or electrical stimuli.

**Fig. 11:**
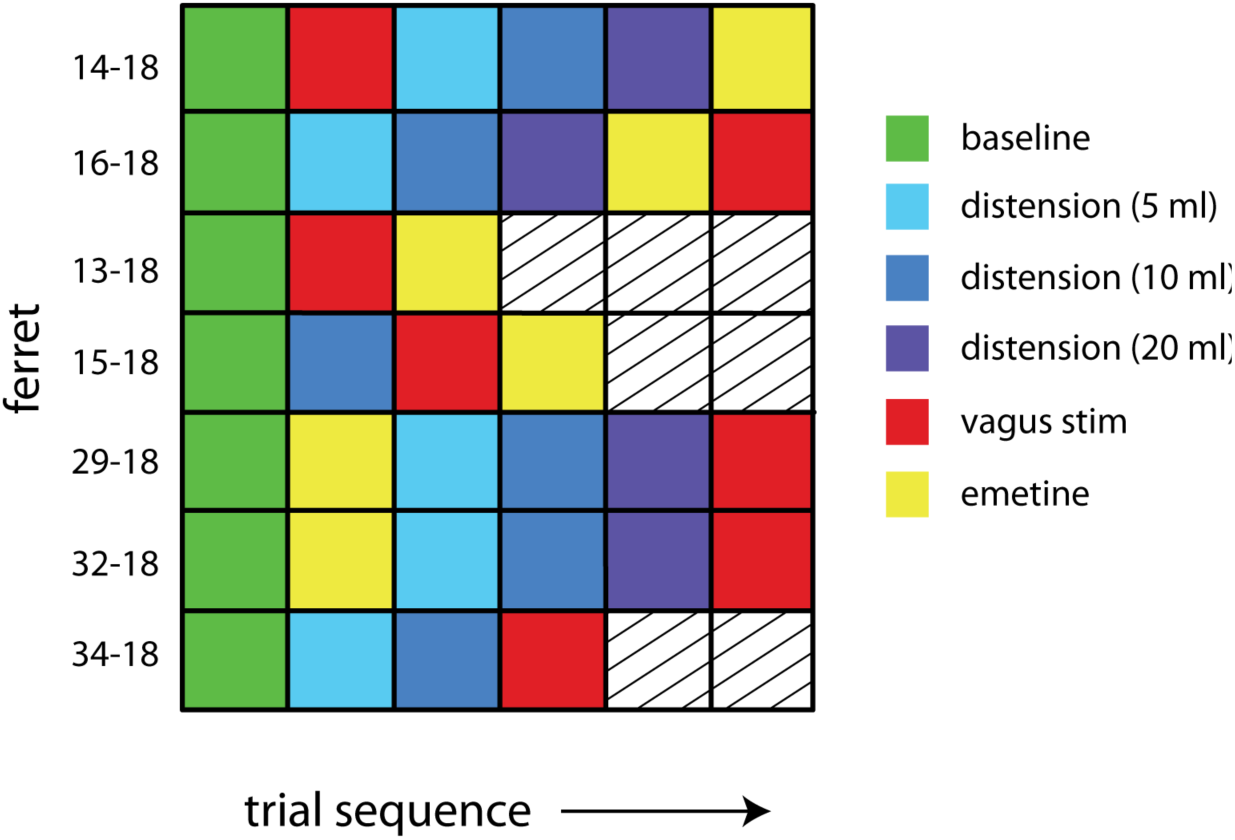
Summary of gastric perturbation trials applied across all acute preparations.

Despite the observed variability in GI signals, standard machine learning algorithms trained on individual subjects were able to detect the state of the stomach with high overall accuracy (Fig. 10).For each animal, the algorithm and the subset of features that resulted in the highest overall accuracy varied. For animals 16-18, 13-18, 14-18, where paddle averaged GI signals were used, band power features alone gave an overall accuracy of 60-70%. Including time series features such as ZX and LL resulted in the overall accuracy values reported in Table 2. Interestingly, for the same 3 animals, GI responses from gastric segment C resulted in the highest accuracy. Additionally, for animal 15-18 where a DF was seen in bipolar paddle averaged responses only, no frequency domain features were required to obtain 81% accuracy. In the context of clinical translation this variability shows that in addition to tuning the parameters of the learning algorithm, individualized feature selection is required to obtain accurate detection of gastric state.

The objective of any closed-loop GI modulation treatment would be to reliably detect the late pre-retch state and deliver an intervention such as electrical stimulation. Therefore, it is necessary that the precision and recall values of the late pre-retch state exceed chance levels. For the 2-state and 3-state confusion matrix precision for the late state is 0.77 and 0.87 and recall is 0.72 and 0.8 respectively. This indicates that the optimal learning algorithms were able to reliably detect the late stage of GI state prior to a retch. Interestingly for the 2-state and 3-state classifier when comparing classifier performance for early versus late stages, the false negative rate was higher than the false positive rate for late stage retch detection (20.39% versus 8.86% and 23.20% versus 13.80% respectively). This implies that the optimal classifier made an incorrect early stage detection more often than an incorrect late stage detection. This result may also be interpreted as the physiological late pre-retch stage starting later than midway between infusion and the first retch as described earlier. For the purposes of this study, the onset of the late stage was arbitrarily set to ensure equal time interval of early and late GI myoelectric signals and was not varied during classifier optimization. It is possible that varying the onset time of the late stage or switching to 2 states for all animals in future studies may lower the false negative rate and improve performance; however, it is also worth evaluating the tolerance for delivering an intervention such as electrical stimulation of the vagus during an erroneous early or late stage detection before optimizing for the false negative rate..

The current investigation is the first to show proof-of-concept for using a machine learning approach to predict GI state. This approach could be applied to treatments of GI diseases and obesity. Indeed, implantable devices are already used to treat these diseases by applying electrical stimulation to the abdominal vagus or gastric surface via continuous or intermittent activation [22, 23], but their efficacy remains questionable [3]. A possible significant improvement of these devices to provide for effective therapy could be the combination of a stimulation approach triggered by monitoring physiological function -- an approach known as closed-loop modulation. Closed-loop devices are being developed for a variety of disorders involving the autonomic nerves and peripheral organs [3]. It should be possible to measure gastric motility with the multi-site electrode and machine learning approach reported here to control electrical stimulation of the vagus to control gastric emptying function, which could affect a change in hunger and satiation; and, therefore limit the control of food intake to treat obesity; a similar approach could be applied to treating GI motility disorders.

## Acknowledgements

This work was supported by NIH funding from the SPARC Program (Award: U18TR002205).

